# *Staphylococcus aureus* gliding comets: formation and observation

**DOI:** 10.1101/2024.12.17.628996

**Authors:** M.C Davies, E.J.G. Pollitt

## Abstract

*Staphylococcus aureus* has been shown to move across soft agar surfaces by several different mechanisms. *S. aureus* can move using either spreading (a form of sliding motility producing generally round/frond like colonies) or by forming comets (slime covered aggregates of cells) that result in long thin dendrites branching out from the central colony. Spreading is agreed to be a form of passive motility whilst the comets are a form of gliding motility (i.e. active). Comets occur under similar conditions to spreading round colonies; however it can be difficult to get the comets to form. Here we examine the variables involved in determining whether comets form as well as report further observations of the comets themselves. We found that the conditions that favoured comet formation (and formed the associated dendrites) occurred over a more limited range than those that enabled spreading motility. Comet formation is very sensitive to the solidifying agents used, the amount of media used and the drying time. We further observed that the comets can propel themselves upwards against gravity unlike spreading motility and that *S. aureus* formed unusual dense aggregates and strand-like structures within comets unlike the normal growing arrangement of *S. aureus* observed in spreading. These results may aid others in producing motility assays to study spreading and comet formation in *S. aureus* and provides further insight into how comets behave.

## Introduction

Historically, *Staphylococcus aureus* was considered to be a non-motile pathogen but it has more recently been shown that *S. aureus* can either move via a passive mechanism called spreading or by forming comets, identified as a form of gliding motility^1–4^. Here we further define the conditions required for comets and for spreading to occur as well as report further observations of how comets are different from spreading and unusual comet behaviours. Kaito et al originally showed that *S. aureus* could expand over surfaces via a passive mechanism called spreading^2^. Spreading is a variation of a type of passive motility called sliding^4^. Sliding occurs in a range of passively motile bacteria. Sliding is a behaviour where bacterial colonies expand outwards as a contiguous monolayer of cells that is physically pushed outwards by growth behind and can be further enabled by surfactant production^5^. In *S. aureus* spreading, the large volume of surfactant and associated moisture initially lifts individual cells and pushes them outwards. As the density of bacteria within the colony increases then this eventually results in the cells at the centre directly physically pushing each other outwards in a manner that is much more like conventional sliding motility^4,6^.

Pollitt et al (2015) previously showed that *S. aureus* can form ‘comets’, which consist of slime-covered aggregates of cells pushing outwards that can leave other *S. aureus* behind^3^. These trails form the pointed dendrites and chains of microcolonies that can be seen around expanding *S. aureus* colonies under the right conditions. These comets have directed movement discrete from the central colony; slime surrounds the moving bacteria, and the comets etch the agar as they move (as well as resisting being washed away by exogenous fluid). Some gliding bacteria, e.g. Pseudanabaena *galeata* and *Isosphaera pallida*, also form ‘comets’ (i.e. defined aggregates that can seed cells behind them as they move)^7,8^. No forms of passive motility have the same characteristics. *S. aureus* comets therefore fulfil the requirements for observationally defining gliding motility as outlined by Henrichsen, however the molecular mechanism required for this behaviour remains unknown^5^. *S. aureus* comets typically occur in assays that use similar formulations to those in which spreading motility occurs so the conditions that favour one type of behaviour remain to be determined. It has been reportedly difficult to generate comets and so the observations were admittedly controversial, however dendrite generation has further been observed with the addition of mucin and a modified assay and this was identified as a separate behaviour from spreading motility^9^. Great variation in motility assays between studies and laboratories is a known problem that leads to misidentifications as has been previously reported with *B. subtilis* and *P. aeruginosa* swarming^10,11^. Consequently this study was aimed at improving and better defining the motility assays in question to aid in further research into *S. aureus* motility^10,12^.

Here we report our investigation into the parameters that affect the *S. aureus* motility assays, to further establish why *S. aureus* moves either by spreading or comet formation, and to understand the conditions under which they occur. We found comet formation were very sensitive to the following: drying times, the amount of media used and the solidifying agent used. Relative moisture available at the surface of the agar is therefore likely critical. We also made new observations of the comets themselves; it was observed that they could move directly upwards against gravity and that the *S. aureus* arranged themselves differently within comets compared to the normal growing arrangement seen in spreading motility. We also did some preliminary studies of the genes required for the two types of motility. These observations provide additional insights into why the *S. aureus* moves either by spreading or comet formation.

## Materials and Methods

### Strains

Cultures of *S. aureus* were grown in Tryptone Soy Broth (Oxoid) for 8 hours at 37°C at 200rpm. The strains used are listed in Table 1. SH1000 WT was used for all experiments, unless otherwise indicated.

**Table 1.** Bacterial strains used in this study.

| Strain | Characteristics | Origin/Reference |
| --- | --- | --- |
| SH1000 WT | Laboratory strain, Repair of 8325-4 strain *rsbU* deletion | O'Neill, A. J. Lett Appl Micro, 2010<sup>35</sup> |
| JE2 WT (USA300) | Laboratory strain, Nebraska library strain | Fey, P. D. et al. mBio, 2013<sup>36</sup> |
| Newman WT | Laboratory strain | Baba et al, J Bacter, 2008<sup>37</sup> |
| SH1000 GFP | Chromosomal GFP | Pollitt et al, Plos Path, 2018<sup>22</sup> |
| SH1000 mCherry | Chromosomal mCherry | Pollitt et al, Plos Path, 2018<sup>22</sup> |
| SH1000 $\Delta pbp3$ | | Simon Foster collection |
| SH1000 $\Delta pbp4$ | | Simon Foster collection |
| SH1000 $\Delta sagB$ | | Simon Foster collection |
| SH1000 $\Delta agr$ | | Simon Foster collection |
| Nebraska Transposon mutant library strains |  | Fey et al, MBio, 2013<sup>36</sup> |

### Motility assay

The motility assay used previously was reused with modifications^3^. The initial standard motility assay used the following: 15 g of Tryptone Soy Broth (Oxoid), 1.25 g of Noble agar (0.25%) (Oxoid) and 460 ml of distilled H_2_0. This was autoclaved, cooled to 50 °C for at least 10 minutes and then used the same day (particular attention was paid to it not being burnt as caramelised sugars inhibit motility). A Glucose solution was also prepared: 40 ml of water and 5 g of D-Glucose (Sigma), which was filter sterilized (0.2 μm filter, Millipore). This was added to the motility media and mixed just before the plates were poured. 25 ml of the combined media was added to 9 cm petri dishes. The plates were left in a laminar flow cabinet with their lids off and upturned next to the plates. The plates were left to dry for 25 minutes. As the drying rate varied according to position in the cabinet, plates were positioned at the back, centre, and front of the cabinet. 5µl of *S. aureus* culture was then spotted at the centre of each plate, the lids replaced, and the plates then incubated at 37°C. The components of this assay have been varied in a systemic way and the results are reported below. All plates were produced as 3 independent replicates unless otherwise stated.

### Variables examined

We examined several different variables, and the details are as follows:

### Solidifying agents and concentration

Bacto agar (Difco, 214010) and Agarose (sigma, A5093) were used as alternatives to Noble agar at the concentrations shown in the Results.

### pH

Subsequent to the addition of Glucose the pH of the media was determined using a SevenEasypH meter (Mettler Toledo) and the pH was adjusted using dropwise addition of either 1M NaOH or 1M HCl.

### Detergents

The detergent Triton X-100 (Sigma, T8787) was tested at the following concentrations: 0.002% (w/v), 0.02 % (w/v), and 0.2% (w/v).

### Observations of the motility plates

We used the following to make observations of the motility plates:

### Light microscopy

A Zeiss Phase contrast microscope was used. Observations were made at 100x magnification.

### Fluorescent Microscopy

An Olympus Epifluorescence Microscope was used to observe motile colonies with either 1% GFP: 99% WT *S. aureus* or 1% GFP: 1%mCherry: 98% WT *S. aureus*. Observations were made with a 10x+10x magnification air lens from above to avoid disturbing the colonies and additional digital zoom. For certain images we used a special 100x MPlan Apo air lens (NA0.95) magnification air lens after we had used a desiccating jar for 30 minutes to remove surface water due to the very short viewing distance). The samples were imaged using a Hamamatsu Orca ER CCD camera using the Volocity software.

### Whole plates

Pictures of the whole plates were taken with the plates suspended on a glass plate and backlit from below with an LED diffuse white light. Black card was placed between the light and the plate to ensure good background contrast^13^. This set up was moved to the warm room for the timelapse video and photos taken every 5 minutes. Images were taken with a Canon point and click camera modified with the Canon Hack Development Kit to enable timelapse photography^14^.

### Nebraska Library screen

The initial motility assay was modified to fit the 120ml square plates with 40ml media per plate, so that multiple isolates from the Nebraska library could be assayed (12 isolates evenly spaced and aliquoted using a 8-channel multichannel pipette to spot every other channel, to ensure enough space to prevent the motile colonies merging). 1µl was spotted for each isolate. The Nebraska library had been prepared as a series of 96 well plates with 200µl of culture for each isolate in each well stored at −80 °C and defrosted as required.

## Results

### Generation of assays that favour or inhibit comet formation

We started the project with our initial assay (see methods, Figure 1). This had been previously reworked from the old motility assay to work in the new laboratory conditions^3^. We again showed that the wildtype (WT) could spread from the inoculation site and produce some dendrites whilst the *agr* mutants moved little from the inoculation site (this was the same for all conditions). This occurs at a lower agar concentration than that shown in Pollitt et al and the dendrites were much shorter than previously reported. At Nottingham University, where the work from Pollitt et al was performed, agar concentration as low as reported here resulted in motility plates that failed to solidify^3^. The assay also produced many more dendrites with SH1000, whereas in Nottingham the Newman strain produced the best-defined dendrites. Given that surface motility assays for other bacteria can vary substantially between laboratories this is sadly unsurprising^10^. From the initial assay, we set out to determine a set of variables that could be adjusted to get 1) an assay protocol where only spreading was present, and 2) an assay protocol that generated as many dendrites (and consequently comets) as possible (examples in Figure 2). We present the variables we studied to obtain an assay that generated well defined dendrites (the dendrite assay) and one that prevented their formation (the spreading assay), and then present the observations made with the dendrite forming assay. We defined well defined dendrites as ones which were over half a centimetre long, were thin and had an obvious tip. If these conditions were met, dendrites generated clear individual comets at their tips as they were forming.

**Figure 1:**
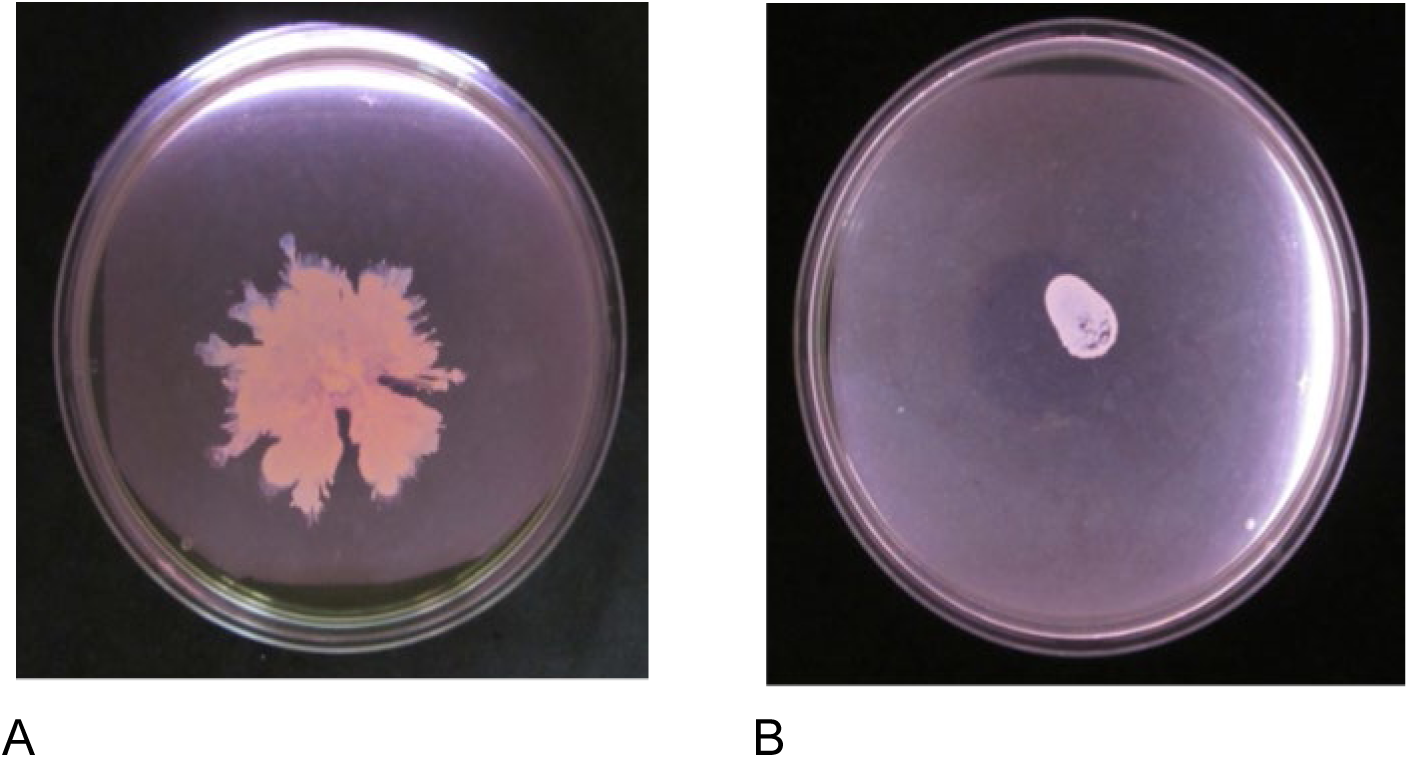
Strains tested under the initial motility assay conditions. A) SH1000 was the most effective strain at spreading and forming comets out of the available wildtype strains with the initial assay. B) The SH1000 *agr* mutant does not move under these conditions and acts as a negative control.

**Figure 2:**
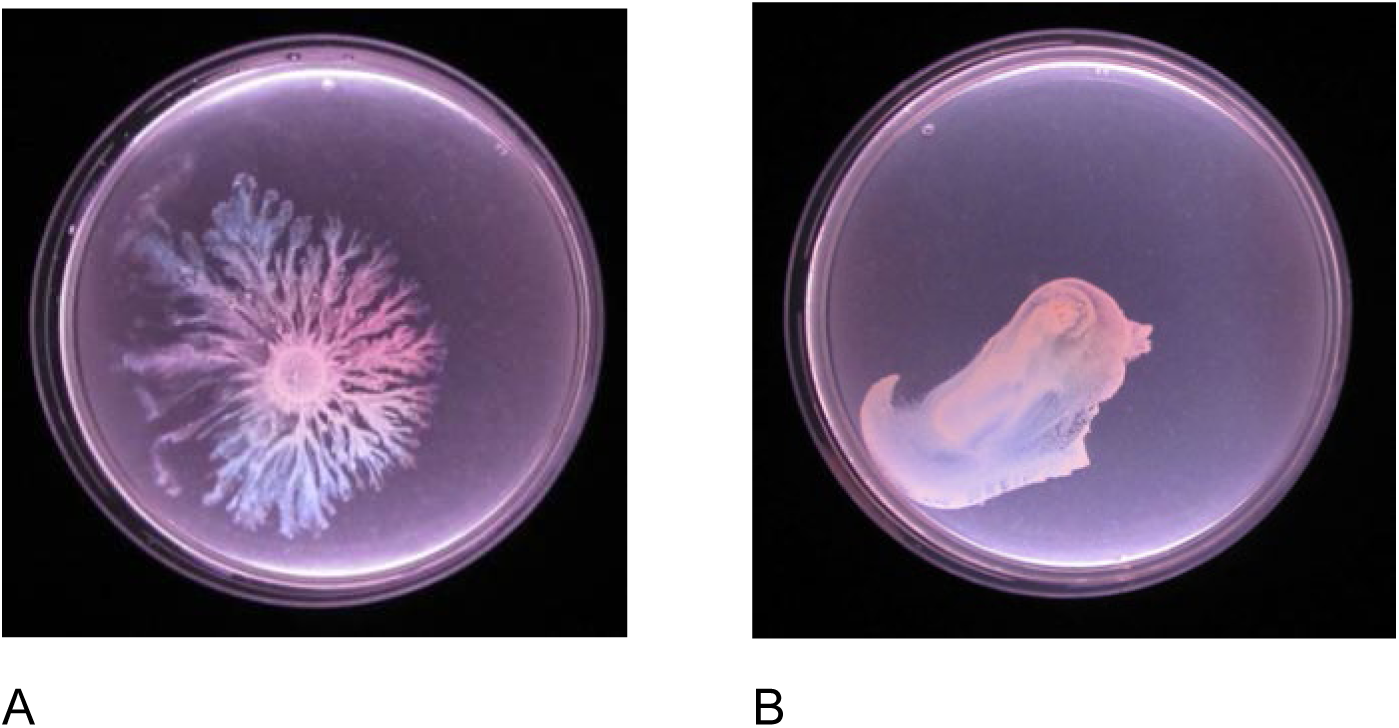
Comet forming and non-comet forming colonies. A) A specimen dendrite forming (where the dendrites are led by comets) colony. B) A specimen non-dendrite forming (spreading only) colony.

### Variables that affected the assay

We examined a range of variables, as outlined in the following sections, which were likely to affect the motility assay and are acknowledged to be important in the design of other motility assays such as the *P. aeruginosa* swarming assay. Our evaluation of solidifying reagents, the effect of variable drying length, pH and the effect of added detergents are reported below.

### Solidifying agents and concentration

We evaluated Bacto agar, Noble agar, and Agarose as solidifying reagents. These have all been used previously in motility assays^3,15,16^. Each solidifying agent was tested in triplicate at concentrations of 0.15%, 0.2%, 0.25%, 0.3%, 0.35%, and 0.5% (Figure 3). 25ml of media was used in each plate. The 0.15% concentration of each agar did not solidify sufficiently to be stable, and disintegrated, with no stable colonies observed. Colonies generated the most dendrites on Noble agar 0.25%, with some restricted movement at higher concentrations. Conversely at 0.20% the entire plate was covered by undefined growth of the *S. aureus*. Bacto-agar produced variable results: the first repeat showed well defined dendrites at 0.2% and some at 0.25%, but some repeats failed to produce any defined dendrites under these conditions. Agarose produced round colonies and, rarely, rounded lobes. Noble agar was used as the final agar of choice in subsequent experiments due to its reliable consistency as a much more purified agar with fewer inhibitory contaminants. At low concentrations Bacto agar varies much more in how well it sets and different batches also were different shades of colour which was a worrying sign and might account for why it worked differently from the work done using Bacto agar previously.^16^.

**Figure 3.**
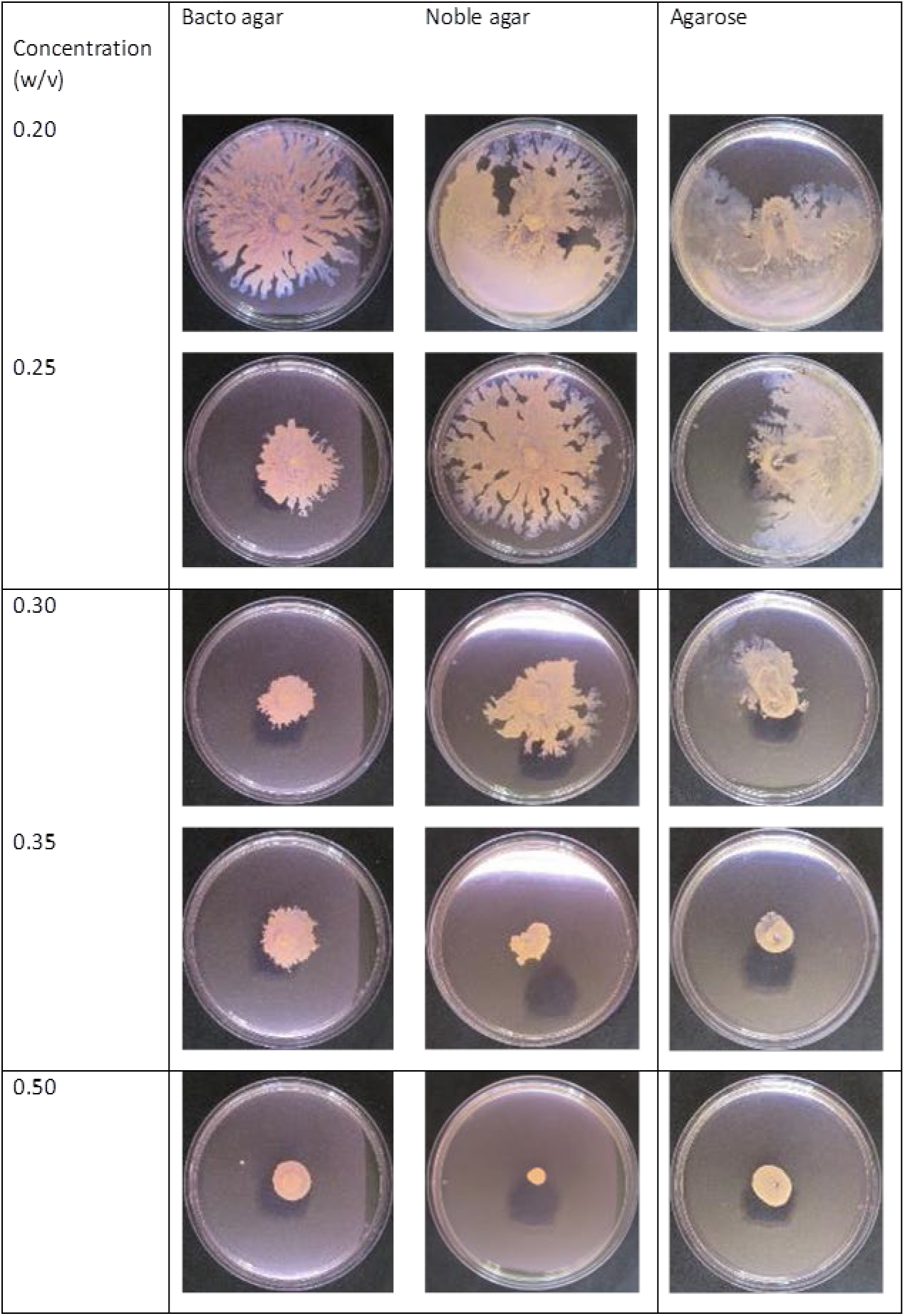
Comparison of the solidifying agents. 25ml of the 3 different solidifying agents (Bacto agar, Nobel agar and agarose) was measured into each plate and left to dry for 25 minutes in the centre of the laminar flow cabinet. The concentration of each type of agar was also varied between 0.2% and 0.5%. As agar concentration increased the dendrite formation and colony expansion broadly decreased. Colonies generated the most dendrites on Noble agar 0.25%.

### Volume and drying times

The moisture content of the media is known to be important in motility assays^10^. Therefore the volume of the media and the drying time, which both influence the moisture content, were studied (using 0.25% Noble agar) (Figure 4). The optimum drying time for 25ml plates was found to be between 25 and 35 minutes; the agar would not solidify if left for less than 15 minutes. We then further tested volumes of 5ml, 10ml, and 15ml for different drying times: 10ml plates appearing to be the most consistent; 5ml plates produced limited numbers of small dendrites; and 15ml plates produced either well defined dendrites or large undefined structures.

**Figure 4.**
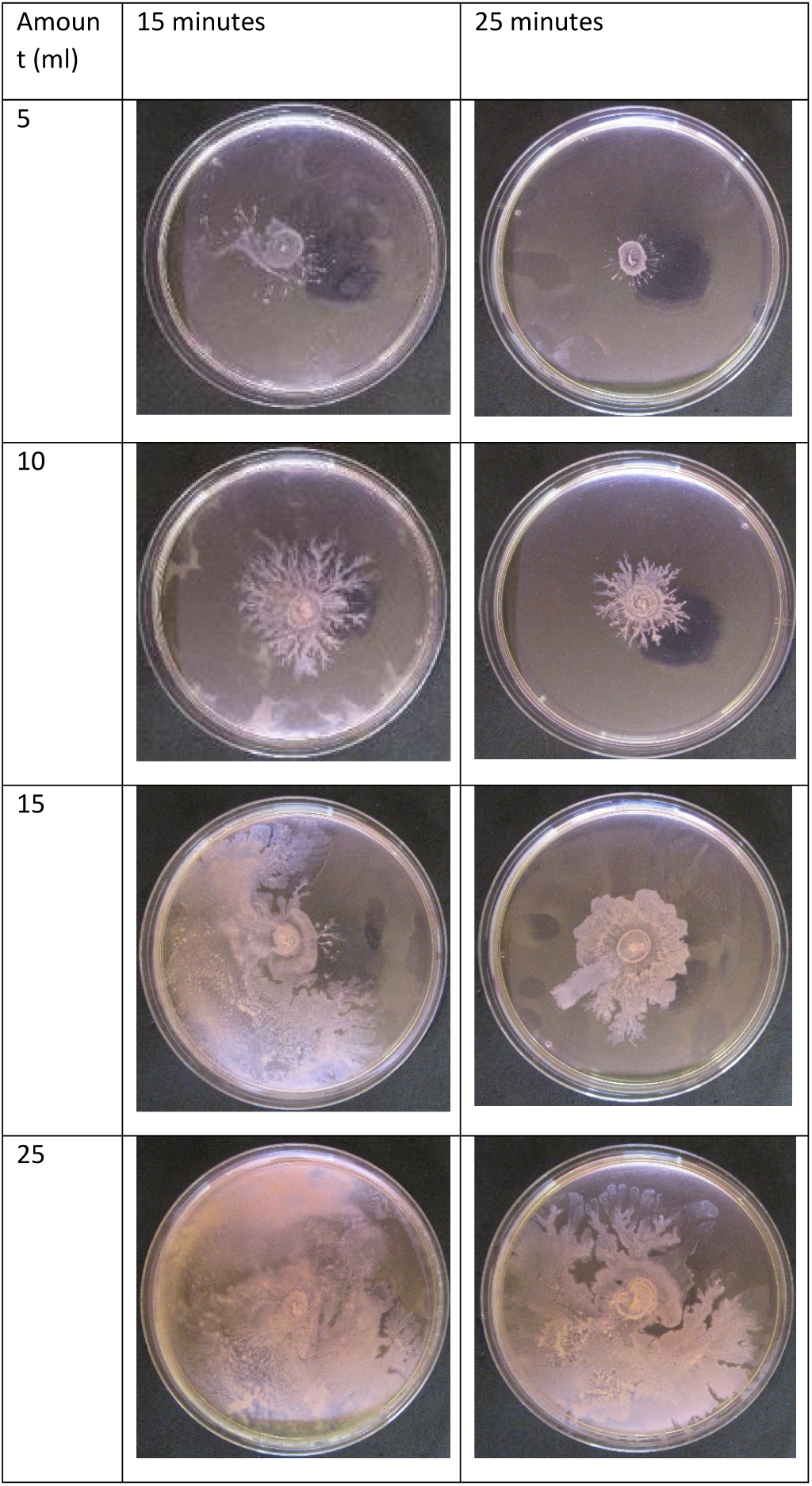
A comparison of noble agar at different volumes and drying times. All plates were made with 0.25% (w/v) noble agar. Combinations of different volumes and drying times were tested. The pointed dendrites associated with comets were more likely to form on smaller amounts of agar. 10ml generated the most consistent dendrite formation.

### pH

Changes in pH were evaluated as it affects motility and growth in other bacteria. Bacterial growth was studied at pH 5.5, 6.5, 7.5, 8.5, and 9.5 (*S. aureus* grows optimally in the range of pH 7-7.5) (Figure 5). pH 7.0 tended to produce the most consistent dendrites, although somewhat lower and higher (6.5-7.5) pH produced similar results. Therefore, the pH was not adjusted further as the natural pH of the media (pH 7) generated optimal results.

**Figure 5.**
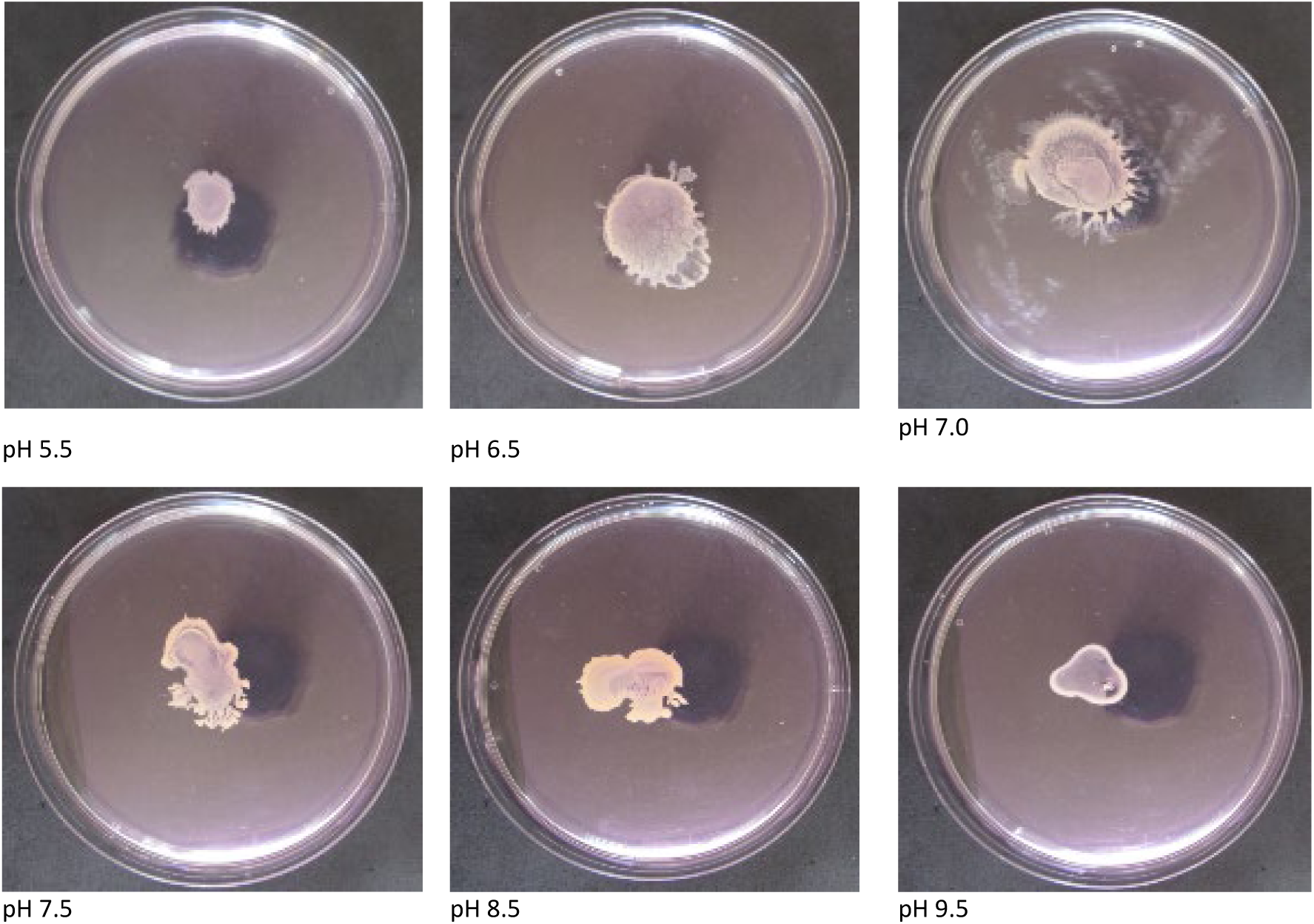
The effect of pH on motility. The pH of the motility media was adjusted with 1M HCl or NaOH as required from the media’s starting pH (6.9). Dendrite formation occurs in a narrower range than spreading alone.

### Incubation with surfactant

Detergents and surfactants can disrupt gliding motility in other bacteria^17^. This can be a problem as trace amounts of washing detergent (detergents generally act as surfactants at low concentrations) can remain in laboratory glassware. We therefore tested the effect of detergent Triton X-100, at 0.002% (w/v), 0.02 % (w/v), and 0.2% (w/v) on bacterial growth (Figure 6). Spreading appeared inhibited at the higher concentration but not at the lower concentration. However in all cases no dendrites were formed.

**Figure 6.**
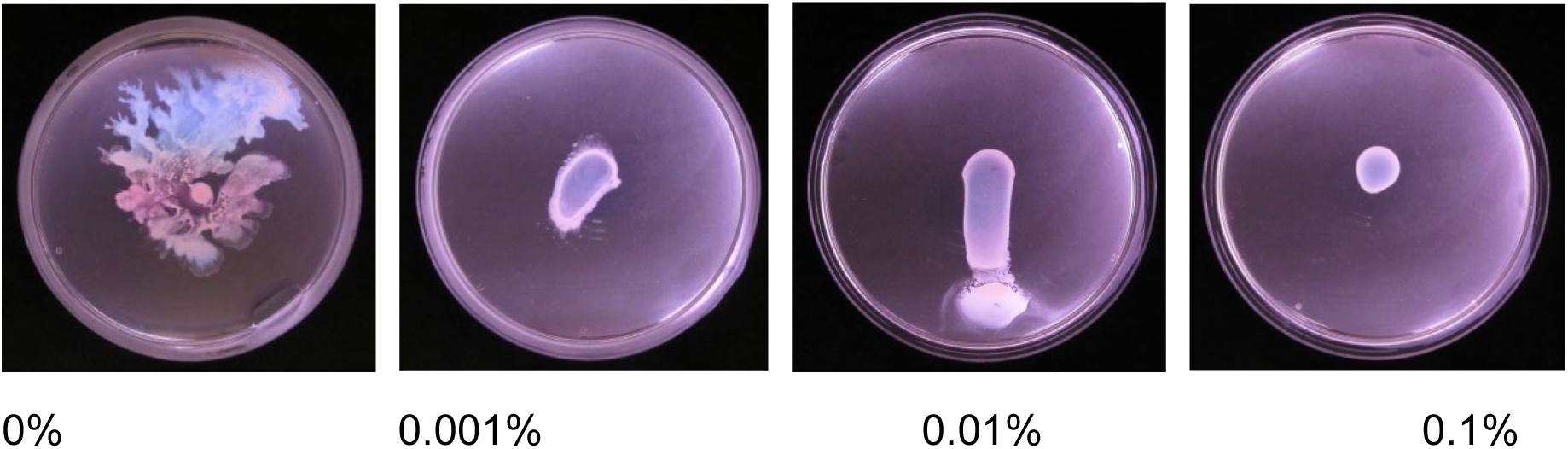
The effect of Triton X-100 on motility. The motility media was adjusted using different amounts of Triton X-100. Increasing amounts reduce dendrite formation and then growth.

### Drying conditions

We observed during the motility experiments that the plates dried at the front of the laminar flow cabinet produced fewer distinct dendrites than the colonies dried at the back (closest to the vent) (Figure 7). There was variation between batches in how large an effect occurred, but the plates at the front generated consistently larger colonies with more spreading. Where possible we prepared two rows of plates (one in the middle and one at the back) so we could obtain colonies that had an acceptable level of colony expansion from the centre. When comparing growth and comet formation we always compared plates from the same row. It was also found that the plates were more consistent and generally produced longer dendrites when they were incubated in an incubator where the temperature and humidity was controlled compared to using a warm room.

**Figure 7.**
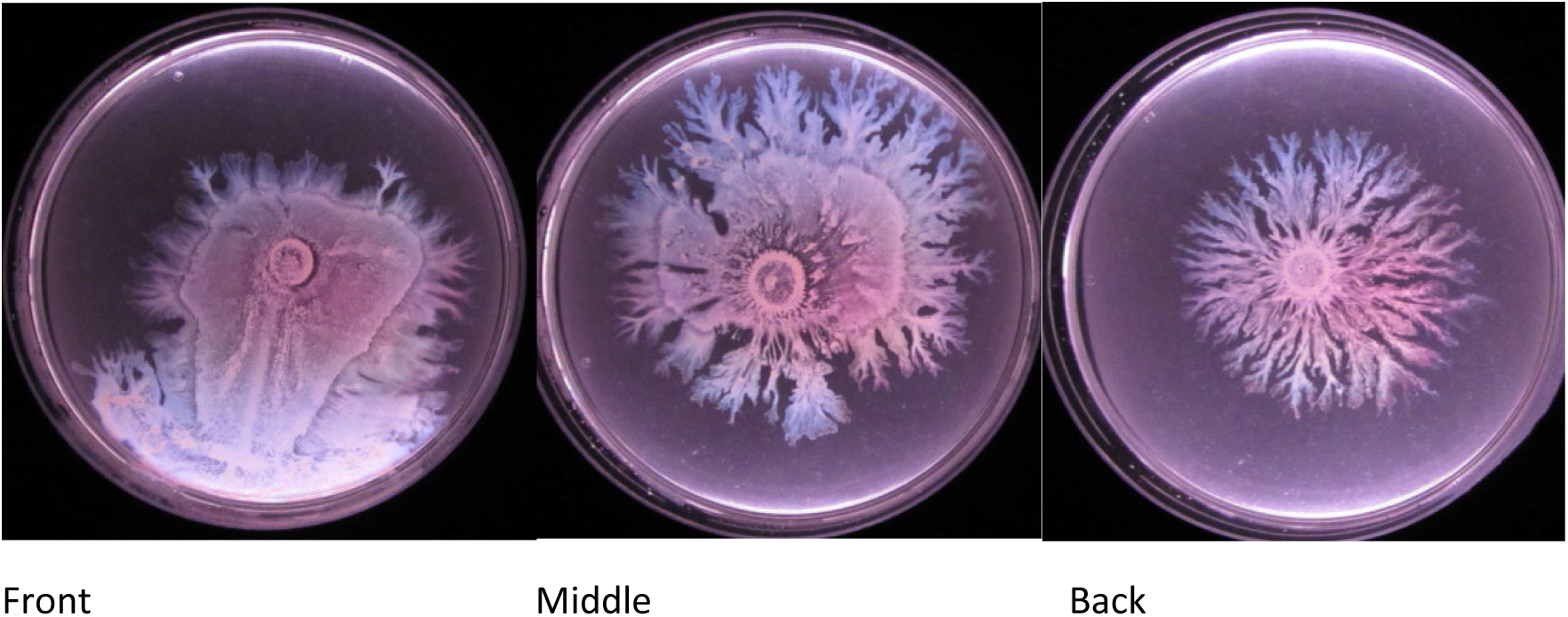
The effect of drying position on the motile colonies. The colonies dried differently depending on where the plates were positioned (Front, Middle and Back of the drying cabinet – back being closest to the air vent). All these plates were generated at the same time and spotted with the same culture of SH1000.

### The optimum assays and evaluation

The optimum finalised assays developed were as follows. The dendrite producing assay used noble agar at 0.25% (w/v), with TSB, water, and purified glucose. 0.625g of Noble agar, 230ml water, 7.5g TSB then autoclaved and 22ml of a stock of 5g of glucose dissolved in 40ml of water (we halved the amount made and filtering is not strictly necessary on a regular basis). 10ml of media was dried as two sets: one for 15 minutes and one for 25 minutes at the back or centre of the laminar flow cabinet. This controlled for variation in the drying rate due to variation in airflow rate in the laminar flow cabinet. The plates were left overnight at 37°C (results appear better in a humidity controlled incubator).

The spreading assay (non-dendrite producing assay) contained 0.5% agarose (w/v)), with TSB, water, and purified glucose. 25ml plates were dried for 25min and then incubated overnight at 37°C.

The dendrite assay was evaluated using a variety of wildtype strains which were spotted on the motility plates and compared (Figure 8). We wanted to determine if the new assay produced more pointed dendrites (and consequently comets) than the old assay when other common strains were used where we had previously demonstrated the occurrence of comets. We found they were being produced well in the new assay with the WT strains tested.

**Figure 8.**
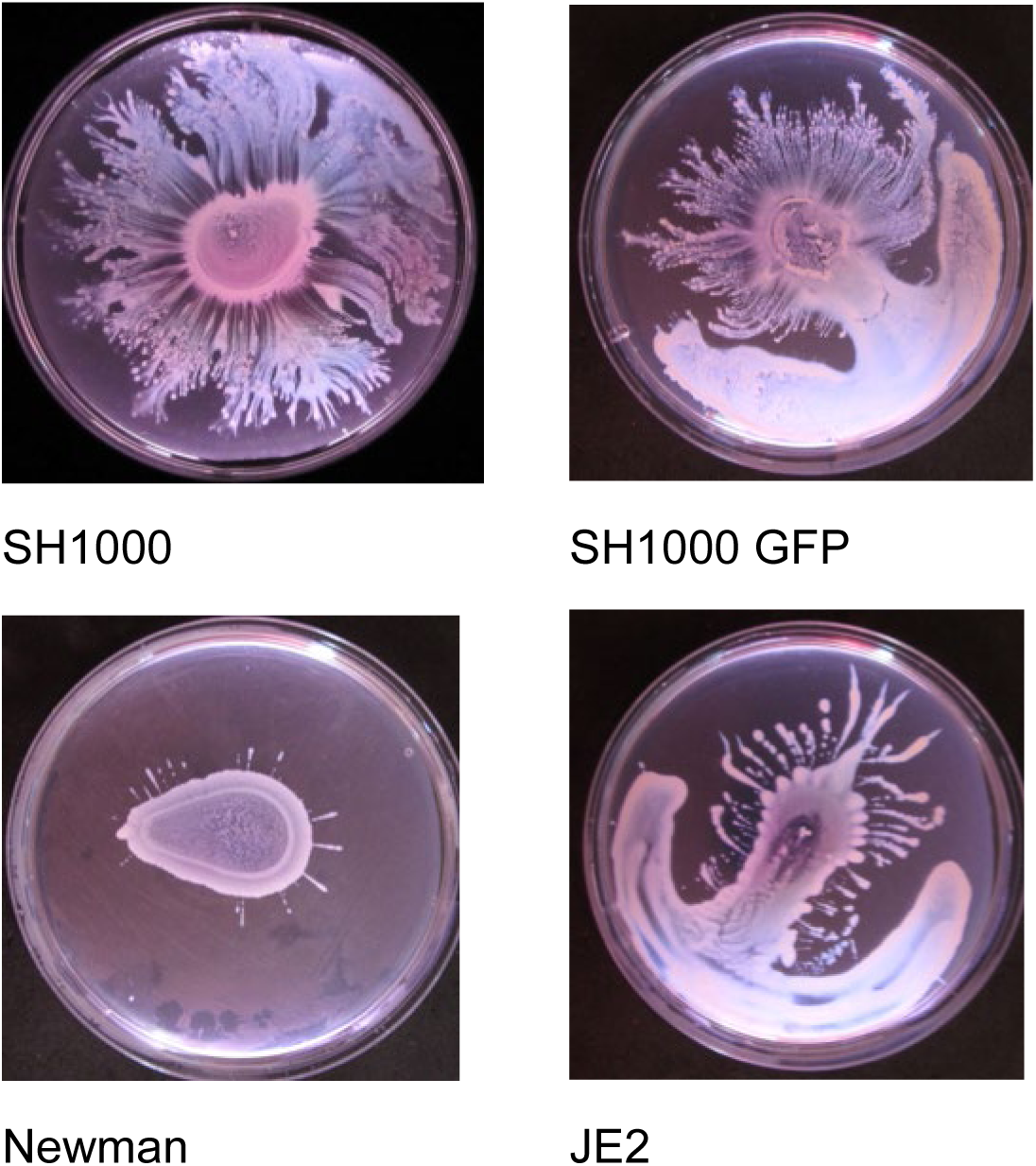
The behaviour of different strains on the finalised media assay. A range of WT strains were tested (SH1000, Newman & JE2(USA300)) and displayed good dendrite formation on the new assay formulation. With the SH1000 GFP and the JE2 plates they were not completely horizontal and so the spreading colony flows downwards to edge and then follows around the edge of the petri dish on the higher amounts of moisture found there.

### Light microscopy

We used a basic phase contrast microscope to confirm visually that comets were still occurring in the dendrite assay (Figure 9). Comet slime tips are easier to see on thinner 10ml agar plates than when they occur on 25ml plates. No slime aggregates were seen in the assay where no dendrites were present. It was also noted that 1) fragments of the slime were also occasionally deposited behind some comets, 2) sometimes so many comets are produced that they produce fans of comets. Where no dendrites were present, dense layers of *S. aureus* were seen resembling previous reports on spreading colonies.

**Figure 9.**
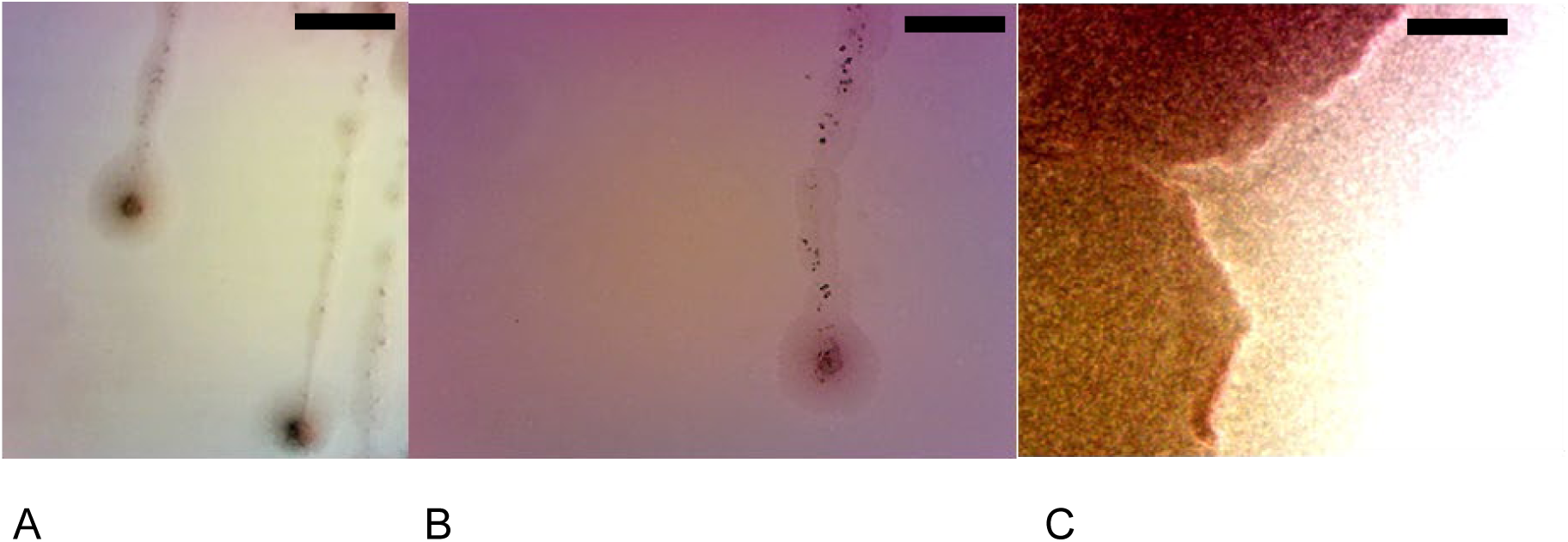
Comets observed using a phase contrast microscope. A) Comets were observed to still precede the dendrites in the finalised media assay. B) Some comets are observed to have trails of detritus behind them along with *S. aureus*. C) Where spreading even distributions of *S. aureus* are seen and no particular structures are observed. Images taken at x100 magnification. Scale bars: 20μm

### Upwards movement

We observed during individual experiments that occasionally when the meniscus of the agar was reached, the colonies could form dendrites (both unbranched and branched) that climbed vertically up the agar against gravity (Figure 10A). To study this observation further with the finalised plate assays we imitated Lin et al and left the plates tilted on the edge of a 96 well plate (Figure 10B)^6,18,19^. Lin et al showed sliding colonies can only flow downwards from the inoculation point with the pull of gravity. Dendrites can however move directly upwards and outwards against gravity; these dendrites being preceded by the comets. Otherwise spreading colonies flow downwards with gravity until they hit moisture collecting at the bottom of the petri dish and then spread out laterally if the plates are sufficiently wet.

**Figure 10.**
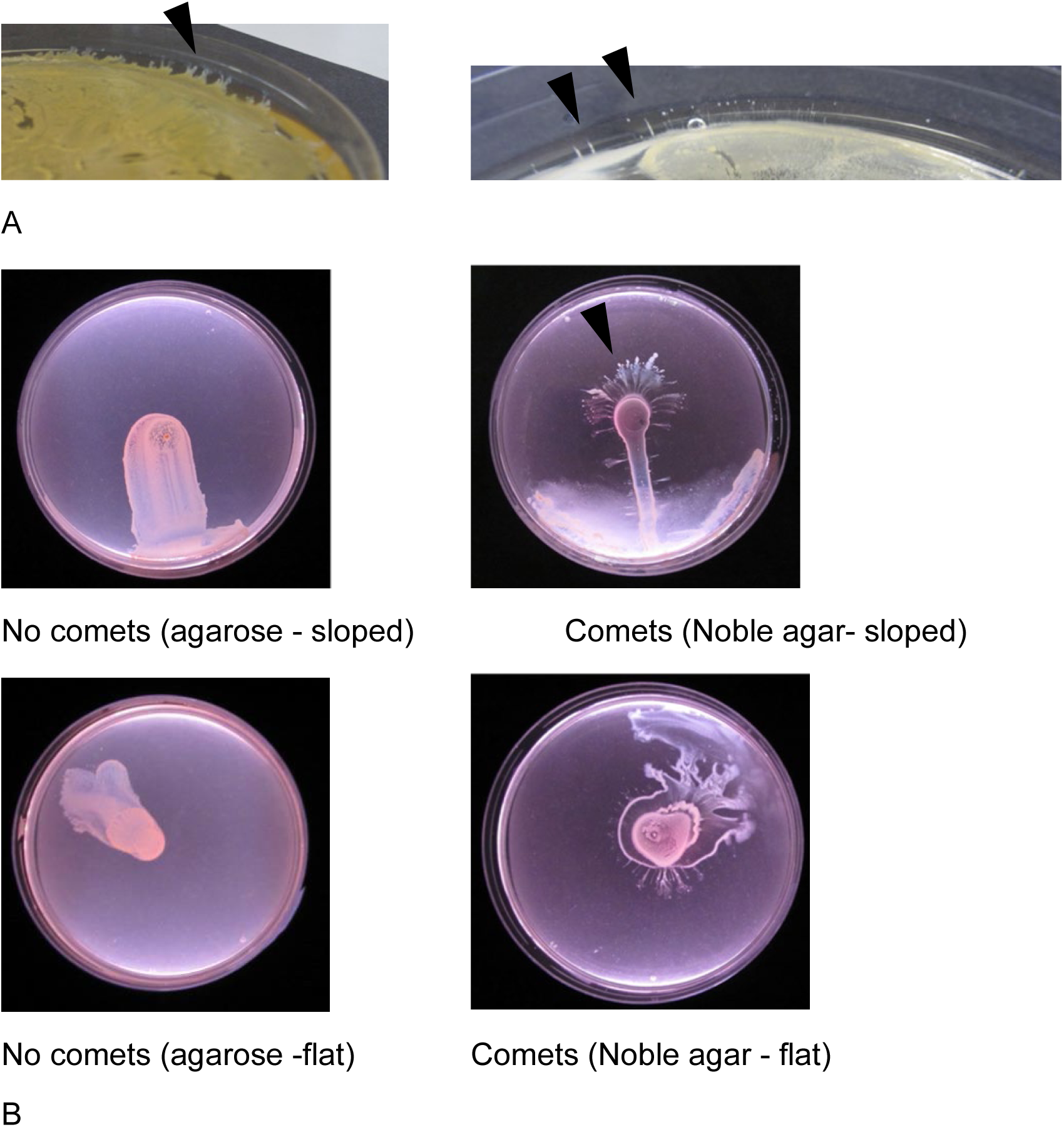
*Staphylococcus aureus* comets can climb surfaces. A) It was noted that when the *Staphylococcus* colonies reached the meniscus, they could directly climb it (Black arrows indicate examples). B) Movement of *S. aureus* on tilted plates comparing dendrite forming and non-forming motile colonies. Dendrites can move directly upwards and outwards against gravity from the inoculation site (see black arrow); these dendrites being preceded by the comets. Otherwise spreading colonies flow downwards with gravity. The controls on a flat surface are also shown.

### The effect of temperature

We also measured the effect of temperature on the comet assay and found that it works over a smaller range than spreading motility (Figure 11). It only works at 33°C and above whilst spreading can occur at 25°C (although extremely reduced). From the way both spreading and comet formation fall off below 37°C, it is important to ensure the plates are incubated in an environment that is maintained at 37°C.

**Figure 11.**
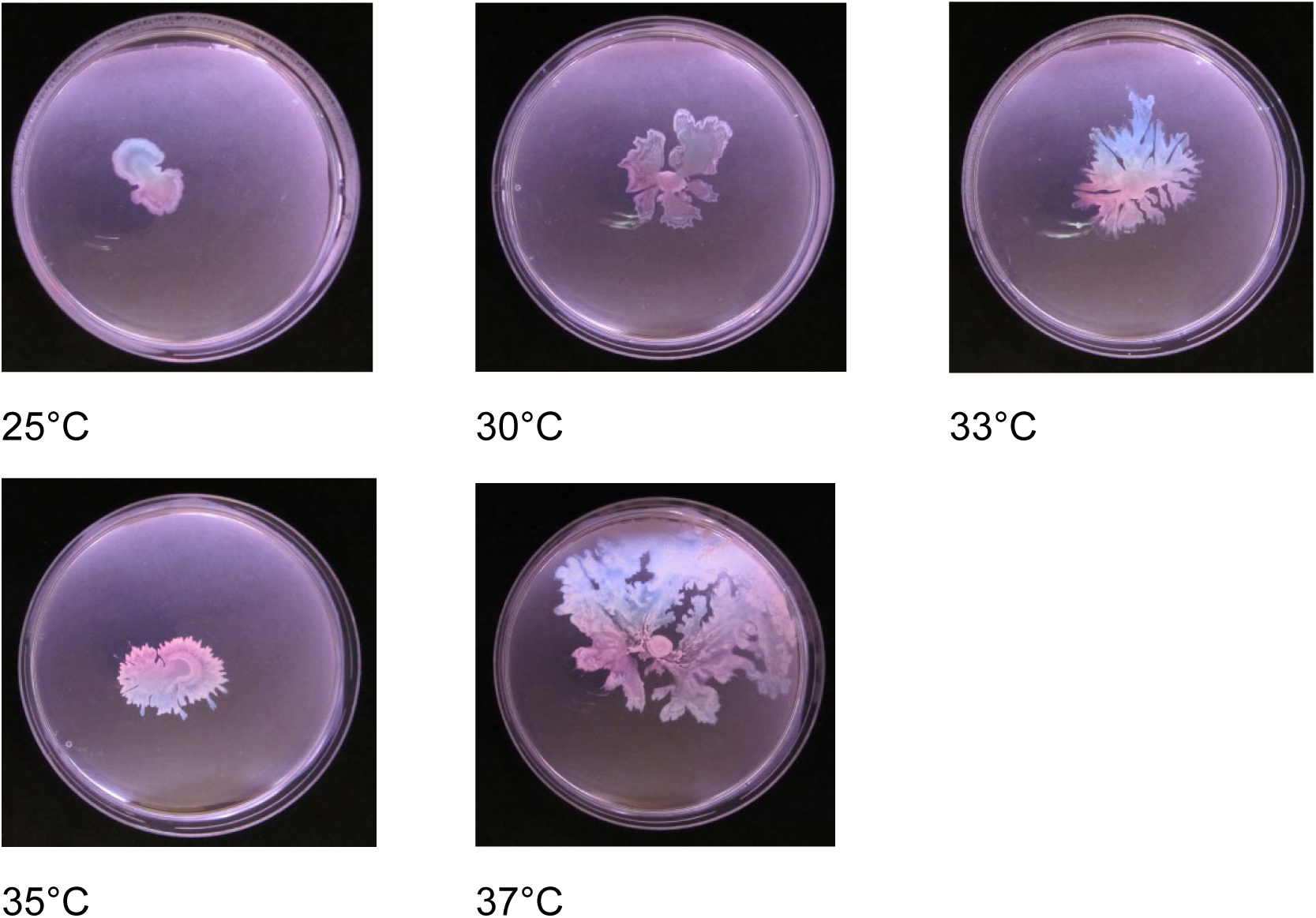
*Staphylococcus* movement at different temperatures. We measured the effect of temperature on the comet assay and found that it works over a smaller range than spreading motility. It only works at 33°C and above whilst spreading can occur at 25°C (although extremely reduced). 37°C is the optimum temperature.

### Fluorescence microscopy

We mixed together low amounts (around 1%) of fluorescent bacteria with WT *S. aureus* (Figure 12). This is a standard motility technique which enabled us to determine the position of individual bacteria within the colony^20^. We found that the fluorescent bacteria were evenly scattered in small groups throughout the expanding colonies (Figure 12D). However, in the same colonies much denser aggregates of cells were formed in the comet tips particularly compared to bacteria in the tail and apparently on the surface of the comets (Figure 12A). We also found that the comet tips was associated with a light diffracting haze, which is likely to be the slime we have noted previously^3^. We have further observed the following: 1) These aggregates tend to form a linear stripe aligned with the direction of the comet (Figure 12A,C), 2) Some comets appear to have multiple strands of bacteria within them coming off a general aggregation of bacteria (Figure 12B), 3) The dense aggregates extend deeper into the agar and further above it than the surrounding bacteria not in a comet tip (Figure 12A), 4) The bacteria do not appear to be fully mobile within the comet as fluorescent bacteria are found the same side of the tail as they are found in the comet(Figure 12B). We already observed previously through phase contrast microscopy that the comets can extend above the agar but had not previously observed them below the plane of the agar surface. This would explain how they can etch tracks in the agar, by physically pushing into it. It would also be a further critical difference to spreading. This work was carried out at x100 magnification. A special x100 air lens (x1000) was not initially usable due to a thin layer of fluid overlying the colony, however by putting the plate in a sealed gas jar with a desiccating agent for 30 min we were able to strip off the surface water. At the higher magnification there was no obvious change to cell shape or other unknown structures (Figure 12E). We also looked at the main body of the colony and found again the bacteria distributed as small groups as per normal *S. aureus* growth (Figure 12F).

**Figure 12.**
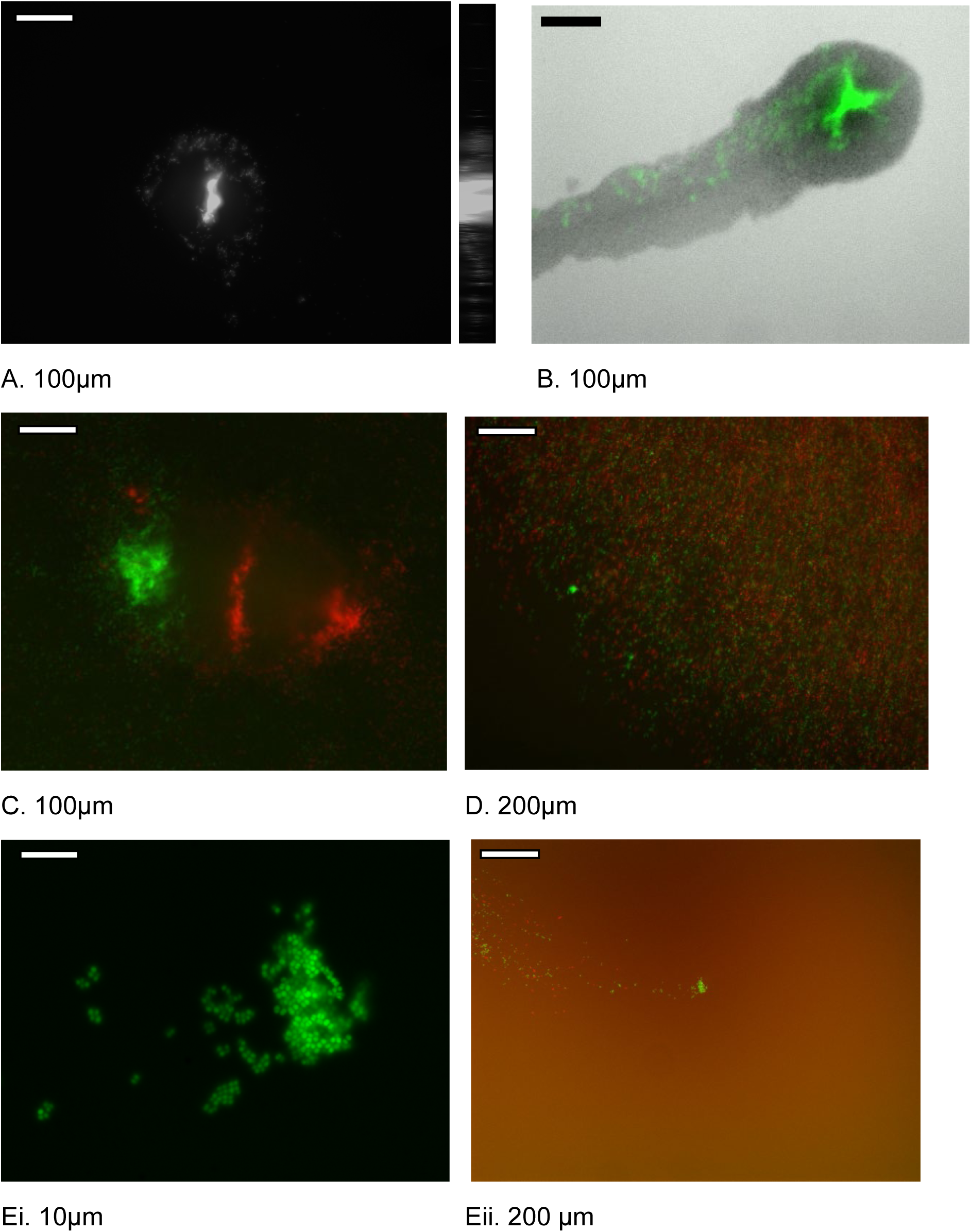
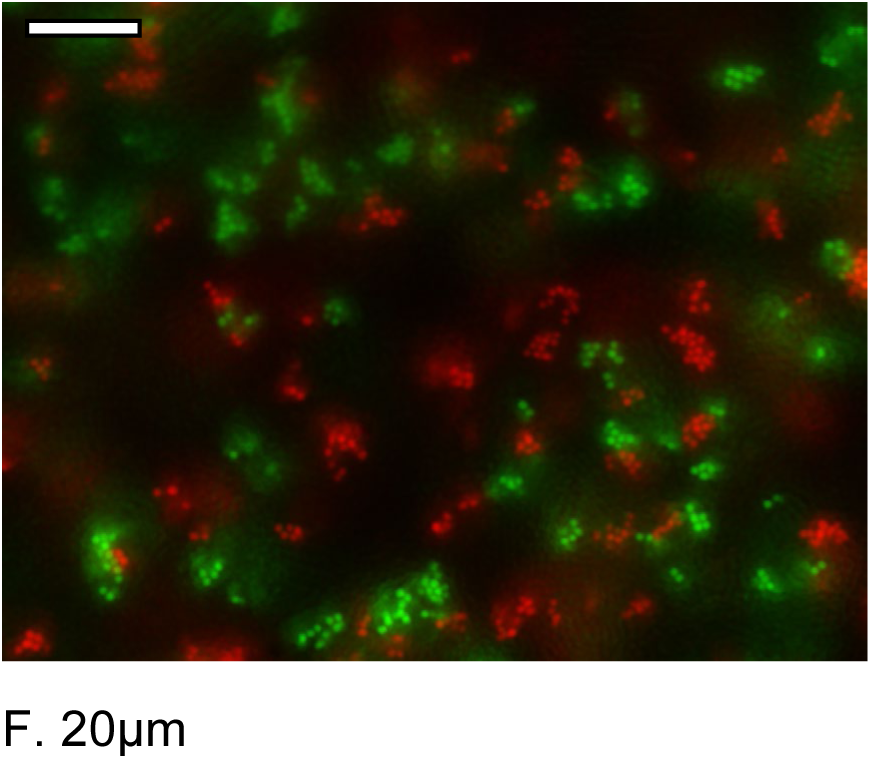
Fluorescence images of *S. aureus* comets. A) Tip of comet from colony with SH1000 1% GFP *S. aureus* and 99% WT, showing the strand of bacteria and haze/slime associated with the comet tip. The cross section is also shown and shows the comet bacteria penetrating below the level of the agar. B) A comet where we also obtained a brightfield (this was rare due to the technical conditions) at the same time. The large aggregate strands are within the comet tip and fluorescence is on one side of the comet only indicating the comet does not dynamically rearrange. C) Comet generated by colony with 1% GFP, 1%mcherry and 98% WT D) Centre of colony with 1% GFP, 1%mcherry and 98% WT. Note the different arrangements to the comets. Ei) x1000 special magnification of Comet generated by colony with 1% GFP, 1%mcherry and 98% WT Eii) x100 overall summary) F) x1000 magnification of centre of the colony with 1% GFP, 1%mCherry and 98% WT. Note even scattering of small clumps as you would expect with *S. aureus* growing normally. Scale bars attached to images.

### Timelapse video

We also made a time-lapse video of the development of a dendrite forming colony (dendrite forming assay incubated in a warm room rather than a humidity controlled incubator due to the constraints of the video equipment). From this video it was observed that at 6 hours the dendrites start emerging from the central colony. This shows there is a step shift from expanding radially to producing comets.

### Identification of candidate mutants involved in comet formation and spreading

J. Tarlit used the initial assay to screen mutants from the Nebraska transposon mutant library^21^. We modified the original assay to fit square plates and found that the JE2 strain did not reliably produce perceptible comets under these conditions (and so M. Davies did the other experiments to look at the variables that were important to create the improved assays). This was not useful for studying comet formation, but it was useful for identifying mutants that did not spread (Figure 13). The following were identified initially as not spreading through two separate repeats, then individual testing and successfully confirmed as not spreading by transduction of the transposon into JE2 WT^22^: Accessory gene regulator protein B (SAUSA300_1989), 5S ribosomal RNA (SAUSA300_0457), Accessory gene regulator protein C (SAUSA300_1991), Lipoate protein ligase A (SAUSA300_1494), ABC transporter, ATP-binding protein (SAUSA300_0978), Accessory gene regulator protein A (SAUSA300_1992), Glycine cleavage system protein H (SAUSA300_0791). One other non-spreading isolate that did not transduce was ABC transporter ATP-binding protein, (SAUSA300_1911). The *agr* mutants act as a good control as they are known not to engage in spreading motility and all *agr* components in the library were detected as non spreaders.

**Figure 13.**
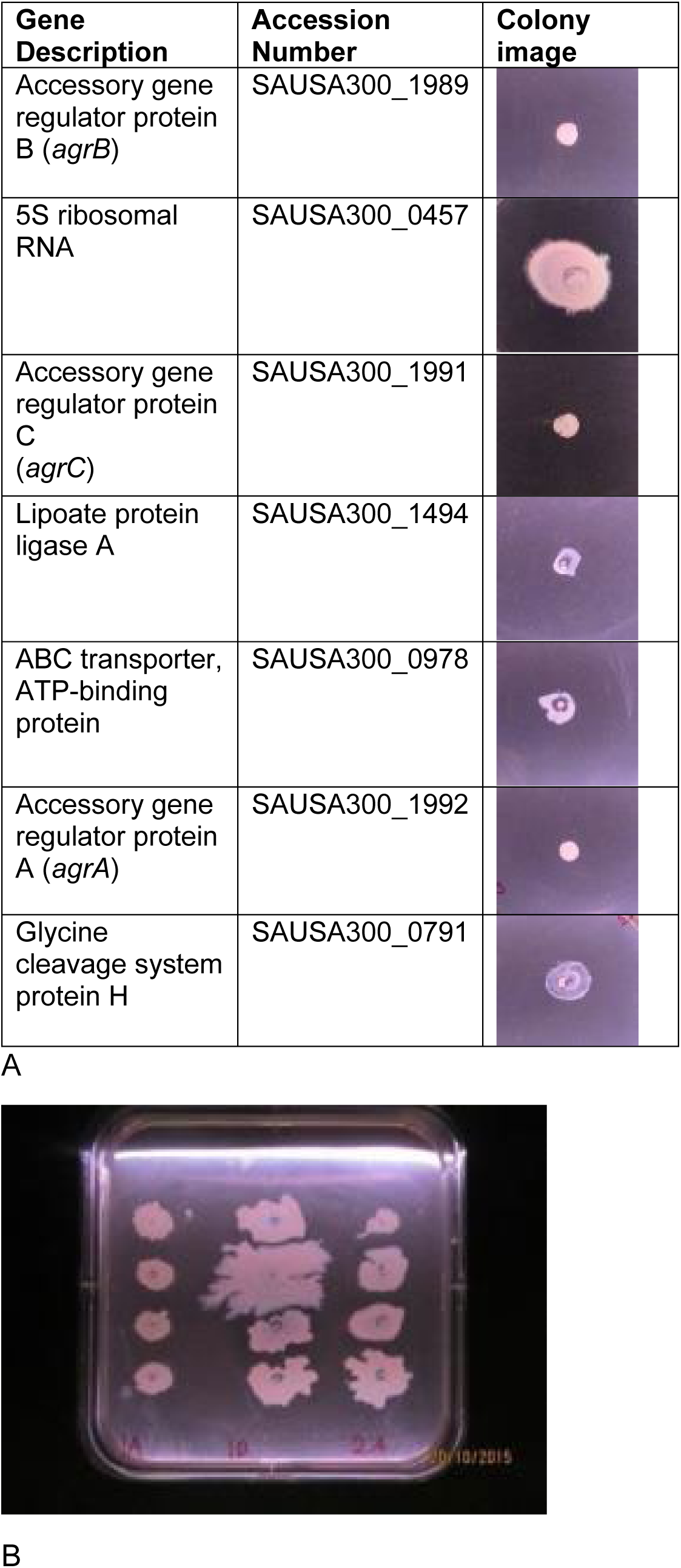
Transposon library analysis. The initial assay to screen mutants from the Nebraska transposon mutant library. A) The spreading colonies of the various identified mutants, the *agr* mutants being previously known. B) An example image of a plate from the screened library spotted with 12 different transposon mutants.

After J. Tarlit had done the work above we continued looking for relevant genes. It had previously been found that *tagO* mutants had eliminated spreading motility (it is more profound than the PSM mutants)^2^. As this is important for the cell wall we decided to investigate other proteins involved in the cell wall, in particular those which are fully conserved, and will grow in liquid media but whose role is relatively undefined. We examined SH1000 *pbp3*, *pbp4* and *sagB* mutants (Figure 14). *Δpbp4* made comets normally whilst *ΔsagB* was deficient and *Δpbp3* was somewhat deficient. However further investigation is required to confirm this.

**Figure 14.**
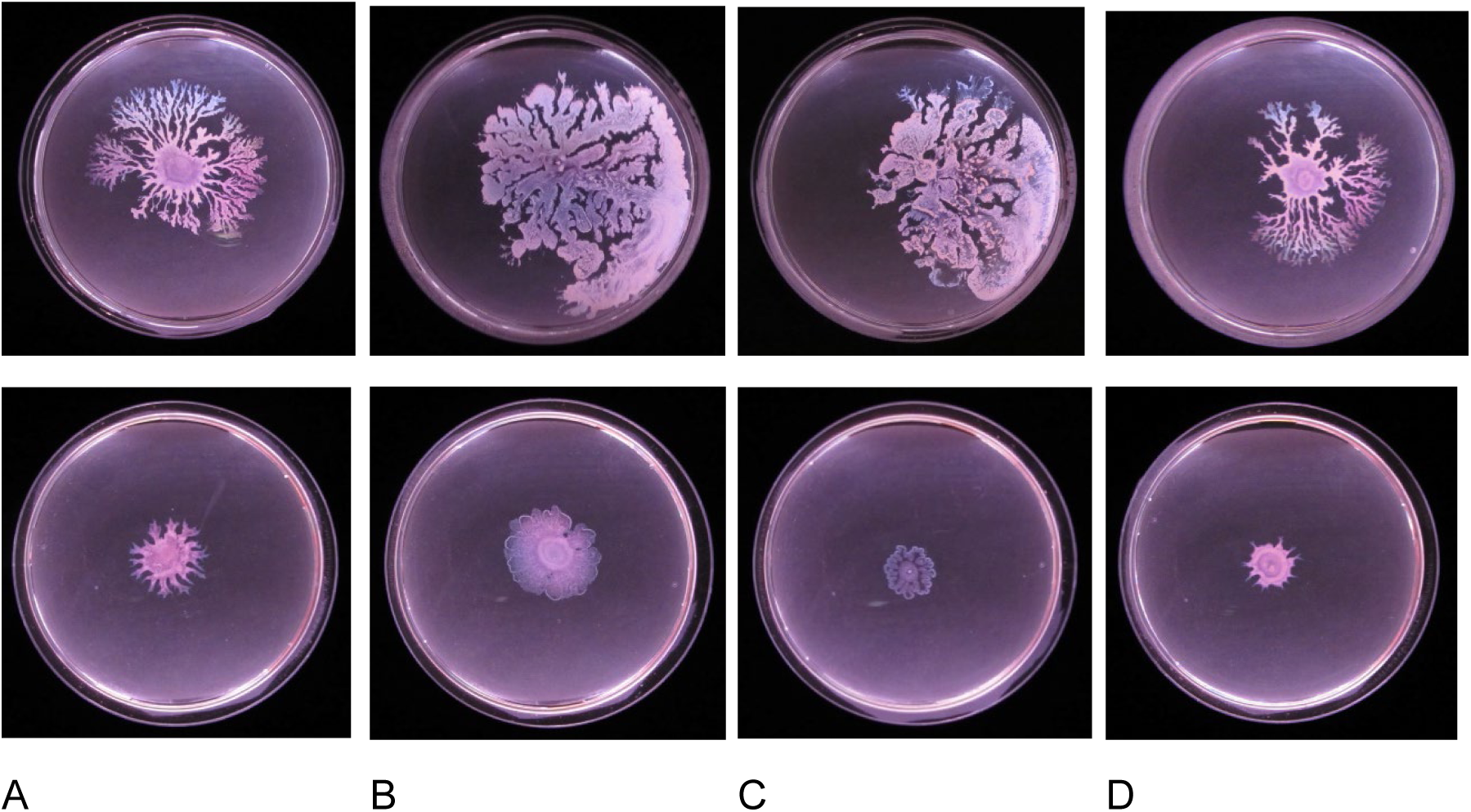
Mutants with altered comet production. A) SH1000 WT control B) SH1000 *pbp3* mutant. C) SH1000 *sagB* mutant. Comets are still produced but these are short and contain a fraction of the mass of WT comets. Comets are completely removed but the colony still spreads, albeit in a ridge like manner. D) SH1000 *pbp4* mutant. This has the same phenotype as the WT. Top row is the 1st batch and bottom row is a second batch where more drying has occurred.

## Discussion

*S. aureus* has previously been found to move over surfaces by two mechanisms: spreading motility and gliding comets^4^. We set out to determine the assay variables which promote motility by spreading alone and which promote and inhibit comet formation. We have developed two assays, one that promotes dendrites formation and therefore the formation of comets (dendrite assay) and another that results in spreading only (spreading assay). Using the new assays we also made further observations that provide an insight into how comets behave, showing their interactions and their ability to climb up surfaces. We have also further studied the genes involved in spreading motility.

### Optimising the assay

The new dendrite forming assay developed in this study resulted in an improvement in the reproducibility and stability of the dendrite-forming colonies we produced. The main differences between the dendrite and spreading assays are that the dendrite forming assay used noble agar vs agarose, the length of drying time and amount of media used. Agarose is believed to be inhibitory in some motility assays as it has a net negative charge compared to agar and may directly interfere with surface binding^16^. This would particularly affect comets as they require attachment to the agar to move, whilst spreading is dependent on the bacteria not being firmly attached to the surface. Moisture is generally required for bacterial movement, however if there is too much moisture in the plate then surfactant and growth alone would be sufficient for movement resulting in spreading and too much surface moisture could interfere with initiation of comet adhesion. The impression is that an only limited range in the water content at the liquid-solid-air interface of the agar (i.e “Wettability”) permits comet formation as it is with other bacteria that are only motile under a limited range of conditions^10,23^. To use the example of bacterial swarming: too much moisture results in swimming motility, whilst too little results in no movement^10^. Swarming occurs at intermediate moisture levels where the bacteria must group together in order to able to move. *S. aureus* comets may be responding to a similar physical situation^12^.

The addition of triton X-100 or a very high pH can prevent the formation of dendrites. It remains to be determined to what extent these directly inhibit growth of bacteria rather than just suppressing dendrite/comet formation alone, but it could be because they interfere with comet formation much more readily than spreading. It appears that comet formation is a much more finely organised behaviour than spreading motility, detergents would particularly disrupt the surfactants being produced by *S. aureus*.

Given the variation in motility assays between different laboratories we recommend that our results should be used as a guide to the variables that affect motility assays from which others researching *S. aureus* motility have the flexibility to develop assays that suit their own laboratory conditions. For instance, a starting point to investigate comets would be using 0.25% Noble agar under a range of drying conditions, using 10ml per plate, ideally in a regulated 37°C incubator with humidity control. Then test a range of drying conditions as we have done. For spreading studies, we recommend using 25ml of 0.5% agarose as a starting point. As it has been found that mucin has a positive effect on dendrite formation it may be a very good research direction to incorporate it or create assays that particularly converge on the conditions in the nose and other locations where *S. aureus* normally colonises^9^. For comets to form there seems to be a general requirement for an energy rich environment that has inherently reduced surface tension or suitable wettability.

### New observations

The motility plates were tilted to observe how the different forms of motility respond to gravity. Dendrites and the associated comets can readily move upwards and outwards from the inoculation site whilst the main mass flowed downwards until it hit the lip of the petri dish and then spread around it (due to presence of water collecting at the bottom of the plate) (Figure 10). The spreading assay resulted in the mass of bacteria only moving downwards until spreading around the moisture deposited at the bottom lip of the plate.

A variation of this assay was previously used to show that spreading was a passive behaviour as the bacteria only flowed down hill and this was attributed to the extraction of water from the substrate and a reduction in surface tension ^6^. As the comets can move uphill and are not carried downhill (unlike spreading) it is further evidence they are actively motile as being able to move against gravity indicates a self-generated force. This result also indicates that *S. aureus* comets require continuous direct physical attachment to the agar substrate in order to move upwards in this way, otherwise they would fall off and flow downwards (and is consistent with track formation)^3^. This attachment was implied previously as exogenous fluid can fail to move the comets whilst sweeping away bacteria in the dendrite left behind ^3^. The effect of gravity on the spreading assay has a notable parallel with how passive motility works in *B. subtilis* left on a tilted surface and is attributed to how the bacteria use surfactant to pull water from the environment and generate a large enough mass to flow downhill^18,19,24^.

We seeded a WT colony with fluorescent bacteria to observe the distribution of individual bacteria within the motile colony (Figure 12). The comets can form dense aggregates of bacteria whereas spreading alone forms diffuse random groupings. The diffuse groupings are explained by surfactant and growth pushing the bacteria outwards radially and breaking them up into small, scattered groups (and is what you expect to see from growing *S. aureus*). Within the comets there are dense aggregates and strands of cells covered with slime (the slime was previously reported but here it is evident as the haze around the comet). The comet structures may arise because the *S. aureus* are not able to dynamically rearrange once they are in the comets (unlike *P. aeruginosa* swarms) and so once the comet forms they stay as an aggregate within a particular location within the comet. This is further supported by the observation that some comets only have fluorescent bacteria on one side of the comet and bacteria were seeded in the dendrite tail only on that side of comet (Figure 12B). The strands of bacteria could be forming due to how the bacteria are constrained by the slime. *Pseudanabaena galeata* also forms non rearranging strands when in its motile comets but its chains of bacteria all point in the same direction^8^. Higher magnification shows that the bacteria do not change shape or local organisation (i.e form chains etc) as some gliding and swarming bacteria do when they move^5,12^. We further see that 1) the slime extends all the way through the comet and 2) comets are embedded into the agar, extending both above and below the *S. aureus* compared to the trail behind (Figure 12). *S. aureus* is unusual in that it is not depositing a slime trail but this can occur in other species^5^. The penetration of the agar likely causes the tracks we have observed forming. Track formation due to partially burrowing in the agar has been observed in other gliding bacteria such as *M. xanthus* but not passively moving bacteria^5,25^.

We took a timelapse video to see if there were any observable macroscopic developments in colony formation. We interestingly saw there was a split between initial expansion and dendrite formation occurring. This also corresponds to the previous observation that comets are not initially present when the bacteria are spotted on the agar but develop later at the edge of the motile colony^3^. This sort of behaviour been seen before with swarming bacteria (particularly *P. aeruginosa*) as they switch from an initial swimming colony moving evenly outwards to the swarming dendrites that move outwards and it takes time to change over(the swarming lag)^12^. In swarming at least this is held to be the junction where the bacteria undergo genetic and phenotypic changes to become swarming proficient. We could plausibly be seeing something similar in that the *S. aureus* need to undergo changes to form the comets.

We also identified candidate genes that affected motility. We focused on identifying these genes using two methods: 1) using the Nebraska transposon library to find defective mutants and 2) testing gene mutants associated with previously reported mutants^21^. We tested all the Nebraska strains with the old motility assay and found a number of Nebraska strains that have defective spreading motility; as an internal control we were able to detect all the *agr* mutants present in the Nebraska library (*agr* mutants do not engage in spreading due to not producing PSMs)^26^. Of the non spreading gene mutants: 5S ribosomal RNA is likely growth defective in these conditions, Lipoate protein ligase A and Glycine cleavage system protein H are part of the Glycine Cleavage Pathway and also likely growth defective in these conditions. SAUSA300_0978 a putative ABC transporter, ATP-binding protein is interesting as it also appears as important for phagocytosis survival. *tagO* is the only non *agr* related spreading mutant previously identified: the *tagO* mutant could be a defective spreader either due to a direct lack of teichoic acids in the cell wall or its growth defect^2^. We therefore looked at other cell wall synthesis genes such as *pbp3*, *pbp4* and *sagB* as they are fully conserved but apparently dispensable for growth in liquid media and we hypothesised they could be more directly involved in motility (Figure 14). All these strains could spread. However, comet formation was almost completely abolished in the *sagB* mutant, and the colony became more frond-like rather than dendritic (the odd plate shows a few comets). Comet formation is also affected in the *pbp3* mutant; comets are still made but they are shorter, and the comet cores were much reduced in size compared to the WT (colony morphology also became frond like). *pbp4* mutants was completely unaffected. *sagB* mutants are particularly known for affecting protein secretion so this may be a factor in the abolition of comet formation. The general lack of genes that affect spreading likely indicates it is purely about PSM surfactant production and growth. It remains to be determined what genes are involved in comet formation.

To restate why we think *S. aureus* comets are actively motile and a form of gliding motility: 1) The comets are directed, discrete movement that resembles known forms of gliding motility in other bacteria 2) The comets produce slime (associated with gliding motility) 3) The comets can produce tracks (associated with gliding motility) 4) The comets are attached to the surface of the agar (Passive motility requires not being particularly attached) 5) Comets can go uphill (indicating self-generated force) 6) Distribution of cells within comets is different from normal growth patterns 7) Comets are not being pushed by the central colony. 8) Comets exhibit specialised behaviours^5,8,17,25,27^. The ultimate question is what is the fundamental mechanism of how the comets move, you can determine if a behaviour is actively motile before the mechanism is determined (as was done for swimming, swarming and twitching motility) but it would increase general confidence if it was found and hopefully this study helps in this direction^5^. If we were to speculate it is possible that the comets could be moving by manipulating the surface tension at the front and back of the comet to dive forward (the tension gradient theory of gliding motility)^28^. Other key areas that need resolution include identifying the composition of the slime. I would not be surprised if it turned out that Phenol Soluble Modulins were involved as amyloid PSM aggregates have been identified in biofilms ^29,30^. It then follows to what extent comet formation overlap with biofilm formation. Sometimes motility reuses biofilm components, sometimes it is antagonistic to biofilm formation^12^. One odd point is how comets can shed bacteria without changing shape or size (in particular, this may involve alternative growth patterns and cell wall modifications as could be involved above).

On a final note when we have looked at protrusions in bacteria we have always found unusual behaviour whether it is *Staphylococcus* or *Mycobacteria*^3,4,16^. We have noted that *Pseudomonas aeruginosa* without flagella and pili can still form swarming like protrusions whilst ‘sliding’ so there may be something interesting happening there too as sliding is only really supposed to produce round and frond-like colonies ^5,31^. Motility mechanisms are regarded as good vaccine targets so unusual ways that bacterial colonies grow and expand should be investigated further to see if they involve novel motility mechanisms^32–34^. It is about finding the conditions where weird phenomena emerge which can then lead to new scientific insights.

### Summary

We have produced two separate assays that allow a split between spreading and comet/dendrite formation and discovered some of the key variables that determine when one form of movement predominates over the other. Based on the dendrite assay we have provided further evidence that *S. aureus* is actively motile as it is capable of directly climbing surfaces and directly observed that the arrangement of individual cells within the comet is different from when the bacteria are spreading. These new observations confirm and extend previous observations of comets made in Pollitt et al^3^. These results and development of better defined assays will aid further analysis of the mechanical basis of comets as well as aiding the detection of mutants that do not form comets.

## Acknowledgements

We would like to acknowledge the support, advice and the use of laboratory equipment of Simon Foster and Egbert Hoiczyk. We would also like to thank Darren Robinson for help with the microscopy. Imaging work was performed at the Wolfson Light Microscopy Facility, using the Olympus Epifluorescence Microscope. We thank Simon Foster for the strains used in this study. M. Davies was funded by a Harry Smith Vacation Studentship awarded by the Microbiology Society, whom we would also like to thank for their generosity. J. Tarlit was funded by University of Sheffield student project fees. I, Eric Pollitt, would also like to thank Dr Stuart Pollitt for editing and comments on earlier drafts of the manuscript.

## Contributions

Conceptualization: E.P. Methodology: M.D., J.T., E.P.; Formal analysis: M.D., J.T., E.P.; Investigation: M.D., J.T., E.P.; Writing - original draft: E.P.; Writing - review & editing: M.D., E.P.; Supervision: E.P.; Project administration: E.P.

## Authorship Notification

Pollitt. E.J.G made repeated attempts to contact John D. Tarlit (J.T.) both by his given permanent email address and available social media but was not able to get a response and thus John Tarlit was unable to review the paper or give permission to submit a preprint to BioRxiv and is therefore not on the author list in accordance with BioRxiv rules. Pollitt. E.J.G supervised and checked the work done by John Tarlit and noted in the text which parts he contributed and is satisfied by his contribution. If he does get in contact I will be happy to add him as an author to reflect his contribution. He did indicate whilst doing the research a few years ago now that he would be happy to have it included in a paper.

---

**Supplementary Video 1 An expanding dendrite forming colony.** A timelapse video of a dendrite forming colony. The video represents 15 hrs of elapsed time. The dendrites emerge 6 hrs after the colony begins to be visible.

